# Spatial and temporal dynamics of SAR11 marine bacteria sampled across a nearshore to offshore transect in the tropical Pacific Ocean

**DOI:** 10.1101/2020.12.31.424995

**Authors:** Sarah J. Tucker, Kelle C. Freel, Elizabeth A. Monaghan, Clarisse E.S. Sullivan, Oscar Ramfelt, Yoshimi M. Rii, Michael S. Rappé

## Abstract

Time-series surveys of microbial communities coupled with contextual measures of the environment provide a useful approach to dissect the factors determining distributions of microorganisms across ecological niches. Here, monthly time-series samples of surface seawater along a transect spanning the nearshore coastal environment within Kāne‘ohe Bay on the island of O‘ahu, Hawai‘i, and the adjacent offshore environment were collected to investigate the diversity and abundance of SAR11 marine bacteria over a two-year time period. Using 16S ribosomal RNA gene amplicon sequencing, the spatiotemporal distributions of major SAR11 subclades and individual amplicon sequence variants (ASVs) were evaluated. On average, 77% of the SAR11 community was compromised of a small number of ASVs (7 of 106 in total), which were ubiquitously distributed across all samples collected from one or both of the end-member environments sampled in this study (coastal or offshore). SAR11 ASVs were more often restricted spatially to coastal or offshore environments (64 of 106 ASVs) than they were shared among coastal, transition, and offshore environments (39 of 106 ASVs). Overall, offshore SAR11 communities contained a higher diversity of SAR11 ASVs than their nearshore counterparts. This study reveals ecological differentiation of SAR11 marine bacteria across a short physiochemical gradient, further increasing our understanding of how SAR11 genetic diversity partitions into distinct ecological units.

## Introduction

The SAR11 order *Pelagibacterales* of the class *Alphaproteobacteria* is one of the most abundant and ubiquitous bacterial lineages on Earth [1]. While found throughout the global ocean and freshwater environments, SAR11 bacteria are particularly abundant in stratified, oligotrophic surface oceans, often making up 25% or more of all bacterioplankton cells [1]. SAR11 are chemoheterotrophic, free-living microorganisms that are uniquely adapted to nutrient-poor environments through small cell sizes and streamlined genomes [2-4]. Similar to other abundant marine bacteria and archaea, most SAR11 lineages are difficult to culture in the laboratory [5], and the current capacity to interrogate SAR11 strains in a controlled laboratory setting remains limited to a small portion of the total SAR11 phylogenetic breadth [6,7].

The SAR11 clade is genetically diverse, with up to 18% small subunit (SSU) rRNA gene sequence divergence distributed among at least five major subgroups [8] and ten subclades including Ia, Ib, Ic, IIa, IIb, IIIa, IIIb, IV, Va, and Vb [9]. Some SAR11 subclades exhibit spatiotemporal distributions that are delineated by depth, season, or geographical location [8,10,11]. At the broad level of subclades, portions of the SAR11 phylogeny appear to differentiate based on ecological parameters, showing attributes similar to bacterial ecotypes. The ecotype concept describes an ecologically homogeneous group of closely related bacteria whose genetic diversity is guided by cohesive forces such as periodic selection, recombination, and genetic drift [12,13]. This concept has been helpful in discerning populations among highly diverse and widely-distributed bacterial groups including *Prochlorococcus* [14] and *Bacillus* [15]. Previous studies of SAR11 have shown evidence for ecotypic differentiation [8,16-18]. Yet, other research has attributed at least a portion of the genetic diversity harbored by the SAR11 lineage to neutral processes [19,20].

The genetic diversity within SAR11 has presented a challenge in understanding the ecological roles and evolutionary origins of the lineage. For example, rather than gene content, measures of SAR11 genomic microdiversity including single-nucleotide polymorphisms and single-amino acid variants were necessary to characterize two ecological niches among closely related populations in the global Ia.3.V subgroup [21]. High intra-population genomic variation has made it difficult to reconstruct SAR11 genomes from metagenomic data [21,22]. Even when SAR11 genomes have been reconstructed from environmental samples, understanding the boundaries between the sympatric populations that they represent has proven difficult [21].

16S rRNA gene sequencing surveys have been foundational in defining SAR11 subclades and examining their spatiotemporal distributions in the environment [8,11,16,23-26]. For example, multi-year studies from the Bermuda Atlantic Time-Series (BATS) in the Sargasso Sea have revealed that the relative abundance of some SAR11 subclades changes with depth and seasonal regimes [11,16]. In other work, a combination of 16S rRNA gene and internal genomic spacer (ITS) analyses were able to discern cold-water (Ia.1) and warm-water (Ia.3) variants of SAR11 subclade Ia [8]. Despite continued efforts to investigate SAR11 bacteria through 16S rRNA gene and ITS surveys, the cultivation of new strains [6,7], metagenomics [18,21,27-29], and single-cell sequencing efforts [30-32], the spatiotemporal characterization of the majority of SAR11 diversity, particularly outside of subclade Ia, remains limited.

One useful approach to investigate the interplay of ecology and evolution within natural microbial communities is to sample across environmental gradients. In the marine environment, light, temperature, pressure, nutrients and other variables form steep gradients with depth in the water column and have been used to investigate the partitioning of microorganisms within different depth horizons of the stratified offshore [33-36]. The nearshore to offshore transition can also offer steep physiochemical gradients in nutrients, salinity, biomass, and productivity which are likely to impact the structure of microbial communities [37]. In tropical coastal environments like those found on islands and atolls, this gradient can be acute; near-island biological, anthropogenic, and physical oceanographic processes provide a substantial source of nutrients for increased biological productivity in otherwise oligotrophic oceanic waters [38].

Kāne‘ohe Bay is a well-studied, semi-enclosed embayment on the windward side of the island of O‘ahu, Hawai‘i [39]. In this study, we used the steep physiochemical gradient provided by the transition from the nearshore environment within Kāne‘ohe Bay to the adjacent offshore environment as a natural laboratory to investigate SAR11 marine bacteria across space and time. This system was sampled monthly over two years to capture the microbial community from ten sites spanning the interior of the bay and the surrounding offshore. We used 16S rRNA gene amplicon sequencing to assess the distribution of SAR11 across a phylogenetic resolution that spanned subclades to individual SAR11 amplicon sequence variants (ASVs). Our findings provide new insight into the ecological differentiation of SAR11 subclades and the distribution of these subclades across the nearshore to offshore continuum.

## Materials and Methods

### Sample collection and environmental parameters

Between August 2017 and June 2019, seawater was collected from a depth of 2 m at ten sites in and around Kāne‘ohe Bay, O‘ahu, Hawai‘i, on a near-monthly basis (20 sampling events over 23 months; **Fig. 1**). At each station, seawater samples for biogeochemical analyses and nucleic acids were collected, and *in situ* measurements of seawater temperature, pH, and salinity were made with a YSI 6600 sonde (YSI Incorporated, Yellow Springs, OH). Approximately 1 L of seawater was prefiltered using an 85 μm Nitex mesh and subsequently collected on a 25-mm diameter 0.1-μm pore-sized polyethersulfone (PES) membrane for nucleic acids (Supor-100, Pall Gelman Inc., Ann Arbor, MI). The filters were submerged in DNA lysis buffer [40,41] and stored at −80°C until extraction.

**Figure 1.**
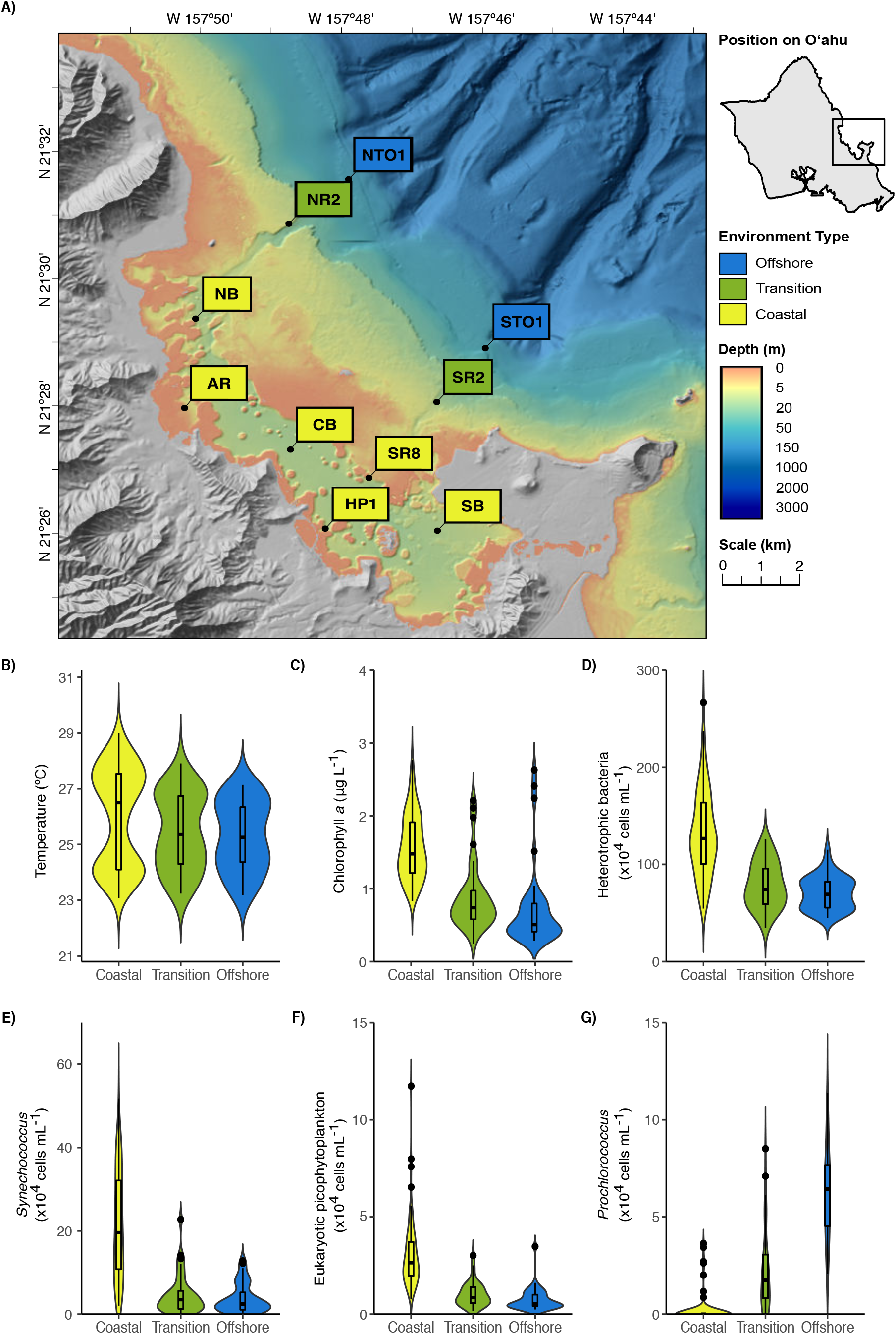
**A)** Map of sampling stations within and immediately adjacent to Kāne‘ohe Bay on the island of O‘ahu, Hawai‘i. Distribution of environmental parameters over two years of sampling across the coastal, transition, and offshore regions of KByT, including **B**) temperature, **C**) chlorophyll *a*, **D**) cellular abundance of heterotrophic bacteria, **E**) cellular abundance of *Synechococcus*, **F**) cellular abundance of eukaryotic picophytoplankton, and **G**) cellular abundance of *Prochlorococcus*. Box plots show the mean and first and third quartile.

Subsamples for chlorophyll *a* were collected by filtering 125 mL of seawater onto 25-mm diameter GF/F glass microfiber filters (Whatman, GE Healthcare Life Sciences, Chicago, IL, USA), and stored in aluminum foil at −80°C. Extraction in 100% acetone and subsequent measurement of fluorescence with a Turner 10AU fluorometer (Turner Designs, Sunnyvale, CA) followed standard techniques [42]. Seawater for cellular enumeration was preserved in 2-mL aliquots in a final concentration of 0.95% (v:v) paraformaldehyde (Electron Microscopy Services, Hatfield, PA) at −80°C until analyzed via flow cytometry. Cellular enumeration of cyanobacterial picophytoplankton (marine *Synechococcus* and *Prochlorococcus*), eukaryotic picophytoplankton, and non-cyanobacterial (presumably heterotrophic) bacteria and archaea (hereafter referred to as heterotrophic bacteria) was performed on an EPICS ALTRA flow cytometer (Beckman Coulter Inc., Brea, CA), following the method of Monger and Landry [43].

Seasons were delineated by fitting a harmonic function to surface seawater temperature collected hourly between 2010-2019 at NOAA station MOKH1 in Kāne‘ohe Bay (https://www.ndbc.noaa.gov/station_page.php?station=mokh1; **Fig. S1A**). Transition seasons (spring and fall) typically experience the greatest amount of change in seawater temperature, thus using the time-derivative of the function thresholds were defined where the derivative was at its greatest absolute value: ≥0.0255 (spring; 30 March through 27 June 2017-19) and ≤-0.0255 (fall; 29 September through 26 December 2017-19; **Fig. S1B**). Summer and winter were defined as the periods between those thresholds (when the derivative was <0.0255 to >-0.0255; summer: 28 June through 28 September 2017-19 and winter: 27 December through 29 March 2018-19).

Using R (version 3.5.1) [44] with the Vegan package [45], stations were grouped into environment types based on non-metric rank-based analysis of a Bray-Curtis transformed distance matrix of environmental parameters (surface seawater temperature, pH, salinity), chlorophyll *a* concentrations, and cellular abundances of *Prochlorococcus, Synechococcus*, eukaryotic picophytoplankton, and heterotrophic bacteria. The pairwiseAdonis package [46] and the ‘betdisper’ function in the Vegan package were used to evaluate the statistical significance and dispersion of these groupings. Comparisons of environmental variables and cellular abundances across the spatial and temporal groupings were conducted using the package multcomp [47] with one-way ANOVAs testing for multiple comparisons with Holm correction and Tukey contrasts.

### DNA extraction and sequencing

Genomic DNA was extracted using a Qiagen Blood and Tissue Kit (Qiagen Inc., Valencia, CA) with modifications [48]. For each sample, 16S rRNA gene fragments were amplified by polymerase chain reaction using barcoded universal primers 515-Y-F and 926R [49]. The 25 µL reactions included 13 µL H2O, 0.5 µL each of forward and reverse primer at 0.2 µM final concentration, 1 µL gDNA (0.5 ng), and 10 µL 1x 5PRIME Hot Master Mix (0.5 U *Taq* DNA polymerase, 45 mM KCl, 2.5 mM Mg^2+^, 200 μM dNTPs) (Quantabio, Beverly, MA, USA). PCR conditions included an initial denaturation at 95 °C for 2 min followed by 30 cycles of 95 °C for 45 s, 50 °C for 45 s, and 68 °C for 90 s, and a final 5 min extension at 68 °C. PCR products were inspected on a 1.5% agarose gel and quantified using the Qubit dsDNA HS Kit (Qubit 2.0, Life Technologies, Foster City, CA, USA). PCR products were normalized to 240 ng each, pooled, and purified using the QIAquick PCR Purification Kit (Qiagen). Pooled libraries were then sequenced on an Illumina MiSeq v2 250 bp paired-end run at the Oregon State University Center for Genome Research & Biocomputing.

### Sequence analysis

The sequence data were demultiplexed and assessed for quality in Qiime2 v 2019.4 [50]. Forward reads were truncated to 245 bp using the --p-trunc-len command in Qiime2 to improve their quality. Reverse reads were not used in these analyses due to poor quality. Sequences were denoised using DADA2 [51] and delineated into amplicon sequence variants (ASVs) that varied by as little as one nucleotide. Denoised sequence data were then assigned taxonomy using SILVA v132 as a reference database [52]. ASVs that contained a minimum of 10 reads in at least two independent samples were retained for subsequent analyses. ASVs undefined at the level of domain and uncharacterized at the level of phylum were excluded from subsequent analyses. Sequences that matched to chloroplasts or mitochondria were included in analyses of community structure but removed in all subsequent analyses of SAR11 including calculations of the relative proportion of SAR11 within the microbial community.

ASVs classified by the SILVA database as the bacterial order SAR11 and the family AEGEAN-169 within the order *Rhodospirillales* (for SAR11 subgroup V) were assigned to SAR11 subclades using phylogenetic placement methods. A 1224 bp alignment of SAR11 reference 16S rRNA gene sequences was created in ARB [53] and used to build a reference tree via RAxML-NG [54]. The SAR11 ASVs identified in this study were aligned with the SAR11 reference alignment and subsequently added to the phylogenetic tree of reference sequences with the EPA-NG algorithm [55], using the best model determined by RAxML-NG for building the reference maximum-likelihood tree [GTR(.960433/4.369000/2.298538/0.794304/6.601978/1.000000)+FU(0.271858/0.192063/0.300 579/0.235499)+G4m(0.276190)]. Visualization of the placement of ASVs on the reference tree used GAPPA [56] **(Fig. S2)**.

Statistical analyses and visualizations were conducted in R packages phyloseq [57], plotly [58], ggplot2 [59], and pheatmap [60], as well as the online tool Venny 2.1 [61]. Generalized linear models (GLMs) were built in mvabund [62] using the ‘manyglm’ function to test for differences in the number of ASVs belonging to SAR11 subclades across seasons and environments. DESeq2 [63] was used to normalize ASV abundance and to evaluate whether individual ASVs exhibited significantly different distributions across seasons or environments using Wald Tests and a false discovery rate of 0.05. The same approach was used to normalize abundance and test for spatiotemporal differences in SAR11 subclades and in total SAR11, *Synechococcus, Prochlorococcus*, and chloroplast abundance.

The frequency of SAR11 ASVs was also assessed by segregating the ASVs into rare (0-5%), low-frequency (>5-25%), mid-frequency (25-75%), and high-frequency (75-100%) categories. Frequency was calculated as the number of samples an ASV was detected in per environment divided by the number of total samples for a given environment (coastal n=120; transition n=40; offshore n=40), multiplied by 100.

Finally, SAR11 ASVs were compared to SAR11 isolates by aligning the SAR11 ASVs to 16S rRNA gene sequences from all known SAR11 isolates using the MUSCLE algorithm in MEGAX [64].

## Results

### Environmental conditions

Non-DNA sequence-based parameters partitioned the KByT sampling sites into three spatial groups, hereafter referred to as coastal (to define the nearshore sampling sites), transition, and offshore (**Fig. S3**). A pairwise-adonis analysis showed statistically significant differences among the three groups (coastal vs. transition: p=0.001, R^2^=0.31; coastal vs. offshore: p=0.001, R^2^=0.43; transition vs. offshore: p=0.012, R^2^=0.071). Dispersion tests found non-significant dispersion (p=0.115, F-value=2.19).

Compared to the transition and offshore stations, coastal stations were defined by lower salinity, higher chlorophyll *a* concentrations, higher cellular abundances of heterotrophic bacteria, *Synechococcus*, and eukaryotic picophytoplankton, and lower cellular abundances of *Prochlorococcus* (**Fig. 1, Tables S1 & S2**). The cellular abundance of *Prochlorococcus* was significantly different across all three environments (p<0.001), ranging from 0.2 ± 0.7 ×10^4^ cells mL^−1^ at coastal stations [mean ± standard deviation (s.d.); n=120] to 2.2 ± 2.1 ×10^4^ cells mL^−1^ at transition stations (n=40) and 6.2 ± 2.3 ×10^4^ cells mL^−1^ at offshore stations (n=40). pH was significantly different between coastal (7.9 ± 0.2; mean ± s.d.; n=120) and offshore stations (8.0 ± 0.2; n=40; p=0.002) and transition (7.9 ± 0.2: n=40) and offshore stations (p=0.015); no significant difference in pH was detected between coastal and transition stations (p=0.841). While not statistically significant, transition stations had lower salinity, higher chlorophyll *a* concentrations, and higher cellular abundances of heterotrophic bacteria, *Synechococcus*, and eukaryotic picophytoplankton in comparison to offshore stations (**Table S2)**. Sea surface water temperature did not statistically differ across the three environments.

An increase in salinity and cellular abundances of heterotrophic bacteria and *Synechococcus* were observed during spring (**Tables S1 & S2**). At coastal stations, salinity ranged from 34.0 ± 1.2 (mean ± s.d.; n=24) during summer to 35.1 ± 1.3 (n=36) during spring, with significant differences between spring vs. summer (p<0.001), fall (p=0.037), and winter (p=0.002). Salinity also reached its highest during spring at transition (35.5 ± 1.2; n=12) and offshore (35.6 ± 1.2; n=12) stations. Increases in the cellular abundance of heterotrophic bacteria between spring and fall were observed in the coastal (p=0.033) and in the offshore environments (p=0.037). The abundance of *Synechococcus* cells increased during spring (25.6 ± 9.7 ×10^4^ cells mL^-1^; n=36) and summer (26.6 ± 12.7×10^4^ cells mL^-1^; n=24) in the coastal environment in comparison to fall (16.8 ± 13.5 ×10^4^ cells mL^-1^; n=24) and winter (16.5 ± 12.×10^4^ cells mL^-1^; n=36; p=0.012). In contrast, the abundance of *Prochlorococcus* cells increased in the coastal environment during fall (0.7 ± 0.1 ×10^4^ cells mL^-1^; n=24). pH, chlorophyll *a* concentration, and the abundance of eukaryotic picophytoplankton cells showed no significant changes over seasons. Surface seawater temperatures were significantly different across all but two seasonal comparisons (spring vs. fall in the coastal environment (p=0.876) and summer vs. fall in the transition and offshore environments (p=0.093 and p=0.157, respectively)).

### Community composition

A total of 2,280 unique ASVs were detected across the 200 samples, including 2,241 bacteria and 39 archaea. The majority of ASVs were distributed across the bacterial phyla *Proteobacteria* (n=1,052), *Bacteriodetes* (n=325), *Verrucomicrobia* (n=60), *Marinimicrobia* (n=54), and *Cyanobacteria* (n=39). A total of 350 ASVs belonged to chloroplasts. A large portion (22.3%; n=509) of ASVs were detected only in the coastal environment, compared to 7.4% unique to the offshore (n=169) and 2.8% unique to transition (n=63) stations.

Measured as a proportion of the entire community, the most abundant orders included the *Synechococcales* (23.3 ± 10.4%; mean ± s.d., n=200), *Pelagibacterales* (21.6 ± 7.5%) *Flavobacteriales* (15.2 ± 7.6%), chloroplasts (11.6 ± 5.6%), *Rhodobacterales* (6.7 ± 3.0%), and *Puniceispirillales* (6.4 ± 4.1%) (**Fig. S4**). Within the *Synechococcales*, the combination of *Synechococcus* and *Prochlorococcus* compromised between 89.0 ± 4.4% and 98.0 ± 2.5% of all the *Synechococcales* relative abundance at a given station. *Synechococcus* dominated the microbial community in the coastal stations (27.4 ± 10.5%, mean ± s.d., n=120), declining in the transition (12.5 ± 8.3%, n=40) and offshore environments (10.7 ± 8.0%, n=40) (**Fig. S5**). In contrast, *Prochlorococcus* was most abundant in the offshore environment (13.3 ± 0.1%), declining in the transition (3.3 ± 0.0%) and coastal environments (0.3 ± 0.0%).

The seasonality of phytoplankton taxa was evaluated using DESeq2 normalized comparisons of chloroplast, *Synechococcus*, and *Prochlorococcus* total abundance as defined by sequence data. No significant seasonal patterns of chloroplast abundance were detected across the coastal, transition, or offshore environments (**Fig S6)**. In the coastal environment, *Synechoccocus* showed increases in spring and summer, and *Prochlorococcus* showed an increase in fall (**Fig S6)**.

### Spatial and temporal distributions of SAR11 subclades

The SAR11 clade accounted for 106 ASVs distributed across eight subclades: Ia, Ib, IIa, IIb, IIIa, IV, Va, and Vb (**Table 1**). Individual samples harbored between 7 and 55 SAR11 ASVs, averaging 14 ± 4 SAR11 ASVs in the coastal environment (mean ± s.d.; n=120), 17 ± 6 in the transition environment (n=40), and 28 ± 9 in the offshore environment (n=40). Generalized linear models with Poisson distributions revealed that the number of SAR11 ASVs recovered per sample varied significantly across environments [Likelihood Ratio Test (LRT)=1168, p=0.001], but did not vary significantly across seasons (LRT=73.49, p=0.114). Of the eight SAR11 subclades identified across KByT, subclade IIa harbored the highest number of unique ASVs (n=41), followed by Ia (n=22) and Ib (n=19) (**Table 1**). Within individual subclades, the average number of ASVs detected in a sample significantly differed between environments for subclades Ib (LRT=641.52, p=0.001), IIa (LRT= 194.92, p=0.001), IIb (LRT= 19.31, p=0.001), and Vb (LRT= 13.78 p=0.001) (**Table 1**).

**Table 1.** Number of SAR11 ASVs detected across KByT.

| SAR11 subclade | SAR11 ASVs (total no.)<sup>a</sup> | Coastal (mean±s.d.)<sup>a</sup> | Transition (mean±s.d.)<sup>a</sup> | Offshore (mean±s.d.)<sup>a</sup> |
| --- | --- | --- | --- | --- |
| Ia | 21 | 5±2 | 4±1 | 4±1 |
| Ib | 20 | 0±1 | 3±3 | 8±3 |
| IIa | 41 | 3±1 | 4±2 | 8±4 |
| IIb | 2 | 0±0 | 0±0 | 0±0 |
| IIIa | 7 | 2±1 | 2±1 | 2±1 |
| IV | 4 | 1±1 | 1±0 | 1±0 |
| Va | 3 | 1±0 | 1±0 | 1±0 |
| Vb | 8 | 2±1 | 1±1 | 3±1 |
| All | 106 | 14±4 | 17±6 | 28±9 |

<sup>a</sup> n=120 coastal, 40 transition, and 40 offshore samples.

The relative abundance of SAR11 subclade Ia was fairly consistent across coastal (14.6 ± 3.9%; mean ± s.d.; n=120), transition (14.2 ± 3.9%; n=40), and offshore (14.3 ± 2.3%; n=40) environments. SAR11 subclade Ia was the most abundant across all stations, ranging from 16.1 ± 3.2% at station SB (n=20) to 13.0 ± 3.6% at station AR (n=20) (**Fig. 2**). Subclade Ib increased in relative abundance in offshore stations compared to their coastal counterparts, exhibiting the highest average relative abundance at offshore station STO1 (10.5 ± 3.7%; mean ± s.d.; n=20) and lowest at coastal station AR (0.03 ± 0.12%; n=20). Similarly, subclade IIa also increased in relative abundance at offshore stations, with the highest average relative abundance at offshore station STO1 (6.3 ± 1.9%; mean ± s.d.; n=20) and lowest at coastal station NB (1.8 ± 0.7%; n=20). On average, subclade IIb was the least abundant of all SAR11 subclades detected throughout KByT, reaching a maximum relative abundance of 0.01 ± 0.04% (mean ± s.d.; n=20) at offshore station NTO1. Collectively, the relative abundance of SAR11 increased from coastal stations (21.6 ± 4.7%; mean ± s.d.; n=120), to transition (23.2 ± 6.4%; n=40) and offshore stations (34.2 ± 7.2%; n=40). DESeq2 normalization of total SAR11 abundance revealed significant differences between the three environments (**Fig. S7)**.

**Figure 2.**
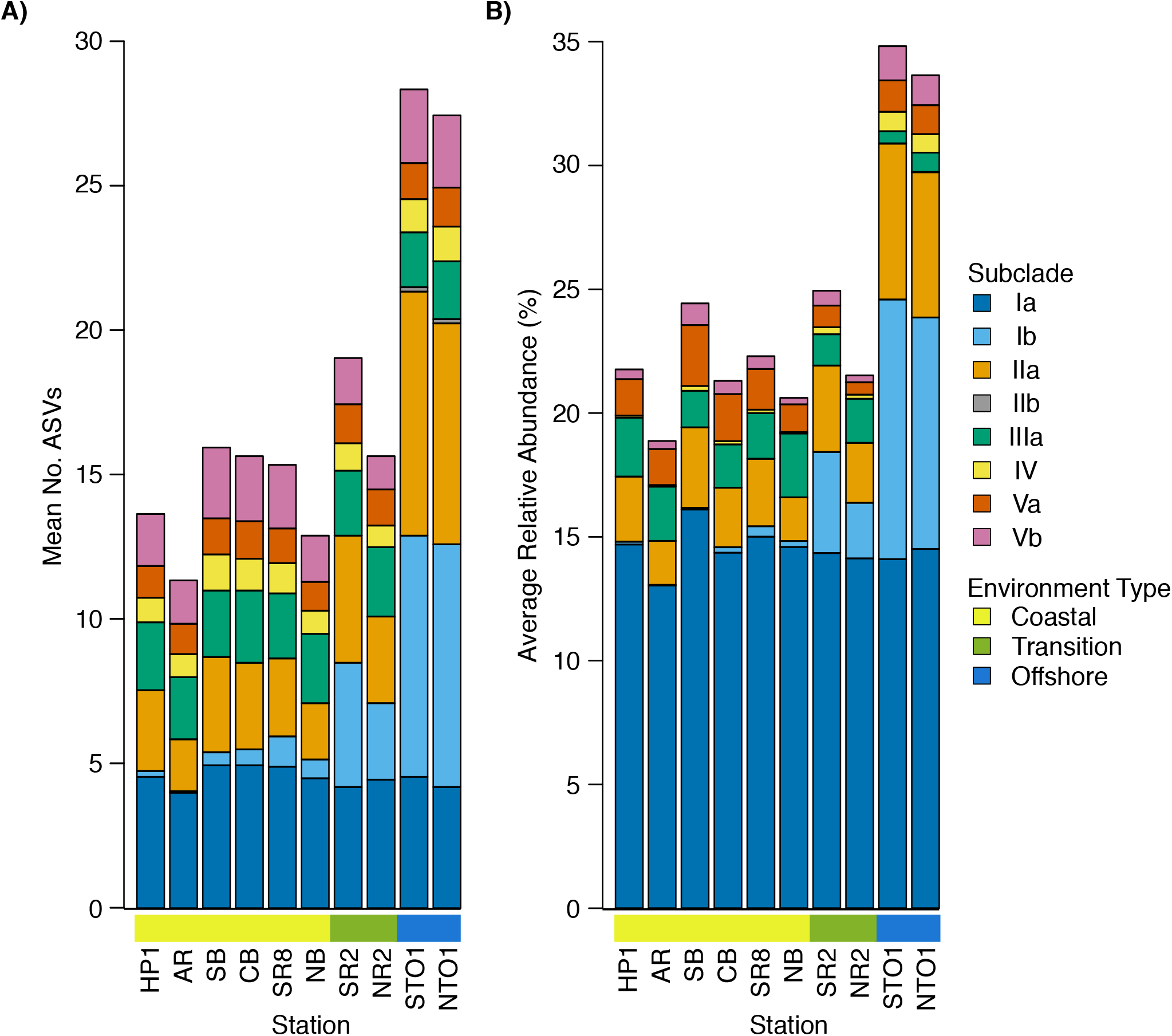
**A**) Average number of SAR11 ASVs and **B**) their relative abundance across KByT stations. n=20 samples per station.

While spatial differences were most pronounced for SAR11 subclades Ib and IIa (**Fig. 2**), significant differences in relative abundance across the coastal to offshore transect were detected for seven of the eight subclades recovered in this study (**Fig. S8**). DESeq2 normalization revealed that subclades Ib, IIa, IV, and Vb were more abundant in offshore compared to coastal stations, while subclades IIIa and Va were more abundant at coastal stations compared to the offshore. Subclades Ia, Ib, IIa, IV, and Va also differed between the coastal and transition environments, while subclades Ia, IIa, IIIa, IV, Va, and Vb differed between transition and offshore environments.

As a whole, the relative abundance of SAR11 did not significantly differ across seasons in the transition or offshore environments but did in the coastal environment between winter and spring (p<0.001) and fall and spring (p<0.001) **(Fig. S7)**. Significant seasonal differences in the relative abundance of individual subclades were observed in both coastal and offshore environments, but not in the transition environment (**Fig. S9**). These include (i) subclade IIIa, which decreased during winter in both the coastal and offshore environments; (ii) subclade Va, which increased in spring for both the coastal and offshore environments; (iii) subclade Ia, which increased during winter in the coastal environment and increased in spring in the offshore environment; (iv) subclade IV, which increased in spring in the coastal environment and increased in winter in the offshore environment; and (v) subclade Vb, which decreased during winter in the coastal environment.

### Spatial and temporal distributions of SAR11 ASVs

Of 106 SAR11 ASVs in total, 39 were found at least once in each of the coastal, transition, and offshore environments (**Fig. 3A**). 20 were unique to the coastal/coastal+transition stations and 44 were unique to the offshore/offshore+transition stations (**Fig. 3A**). All subclades except IIb, which was exceedingly rare overall, contained differentially distributed ASVs across the three environments when evaluated by DESeq2 normalization (**Fig. S10; Table S3)**.

**Figure 3.**
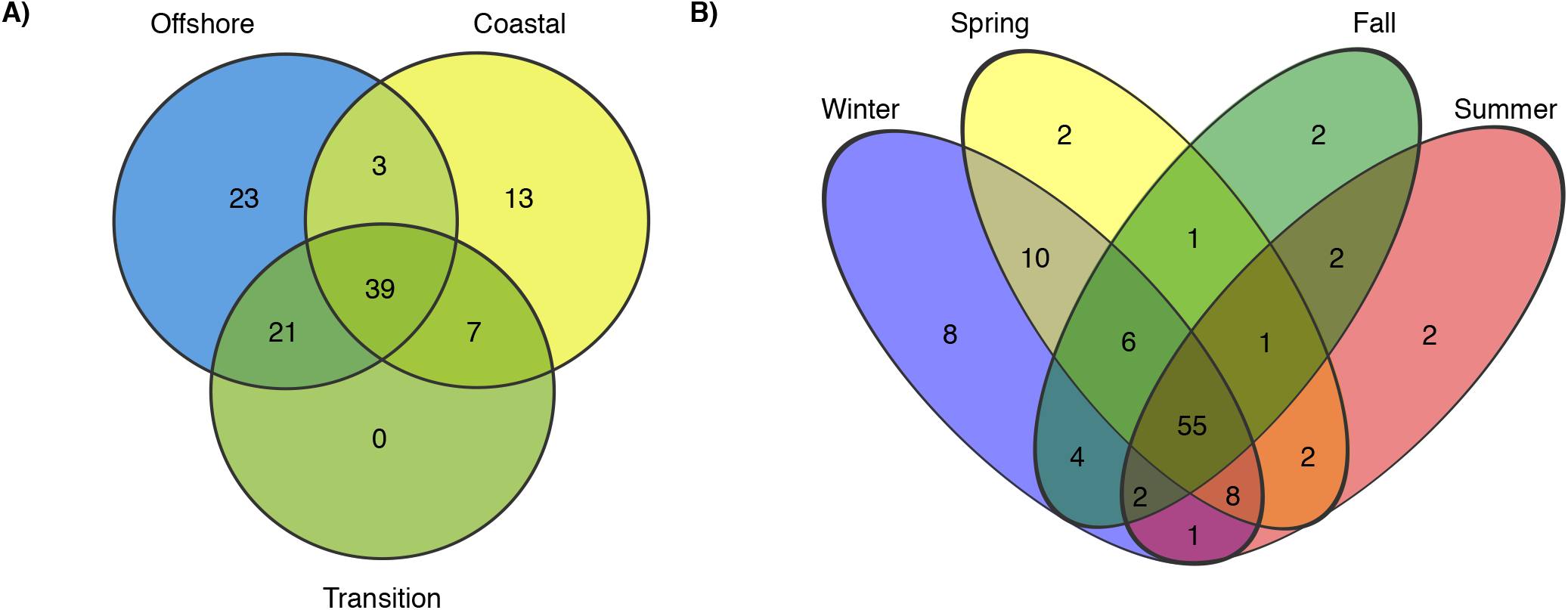
Venn diagrams comparing the distribution of total number of ASVs detected across **A**) environments and **B**) seasons.

Over half (55) of the 106 SAR11 ASVs appeared across all four seasons (**Fig. 3B**). The fewest number of SAR11 ASVs were detected in the fall and summer (73 ASVs and n=40 samples each), while a higher number of SAR11 ASVs were detected during spring (85 ASVs in n=60 samples) and winter (94 ASVs in n=60 samples). Some ASVs (n=14) were restricted to a single season: two in each of spring, summer, and fall, and eight in winter. All of the season-specific ASVs also had restricted spatial distributions, were infrequently recovered (n=2-5 samples), and were relatively low in abundance (<1.5% of the SAR11 community). Using DESeq2 normalization, 18 ASVs showed seasonal differences in at least one of the three environments (**Fig. S10; Table S3**). While 11 of these seasonal ASVs were found in the coastal environment, four were detected in the transition environment, and only two were recovered in the offshore environment.

### Frequency of SAR11 ASVs across KByT

SAR11 ASVs were grouped into four categories based on the frequency they were detected in samples within each of the three environments: rare (<5%), low-frequency (5-25%); mid-frequency (25-75%); and high-frequency (>75%). In general, most SAR11 ASV diversity was comprised of either rare or low-frequency ASVs, including 45 of 62 ASVs in the coastal environment (73%), 47 of 67 ASVs from the transition environment (70%), and 52 of 86 ASVs from the offshore environment (60%) (**Fig. 4**). The offshore environment harbored 16 ASVs in the high-frequency category, followed by nine in the transition environment and seven in the coastal environment. Of these, only three were high-frequency across all three environments and none were unique to the transition zone.

**Figure 4.**
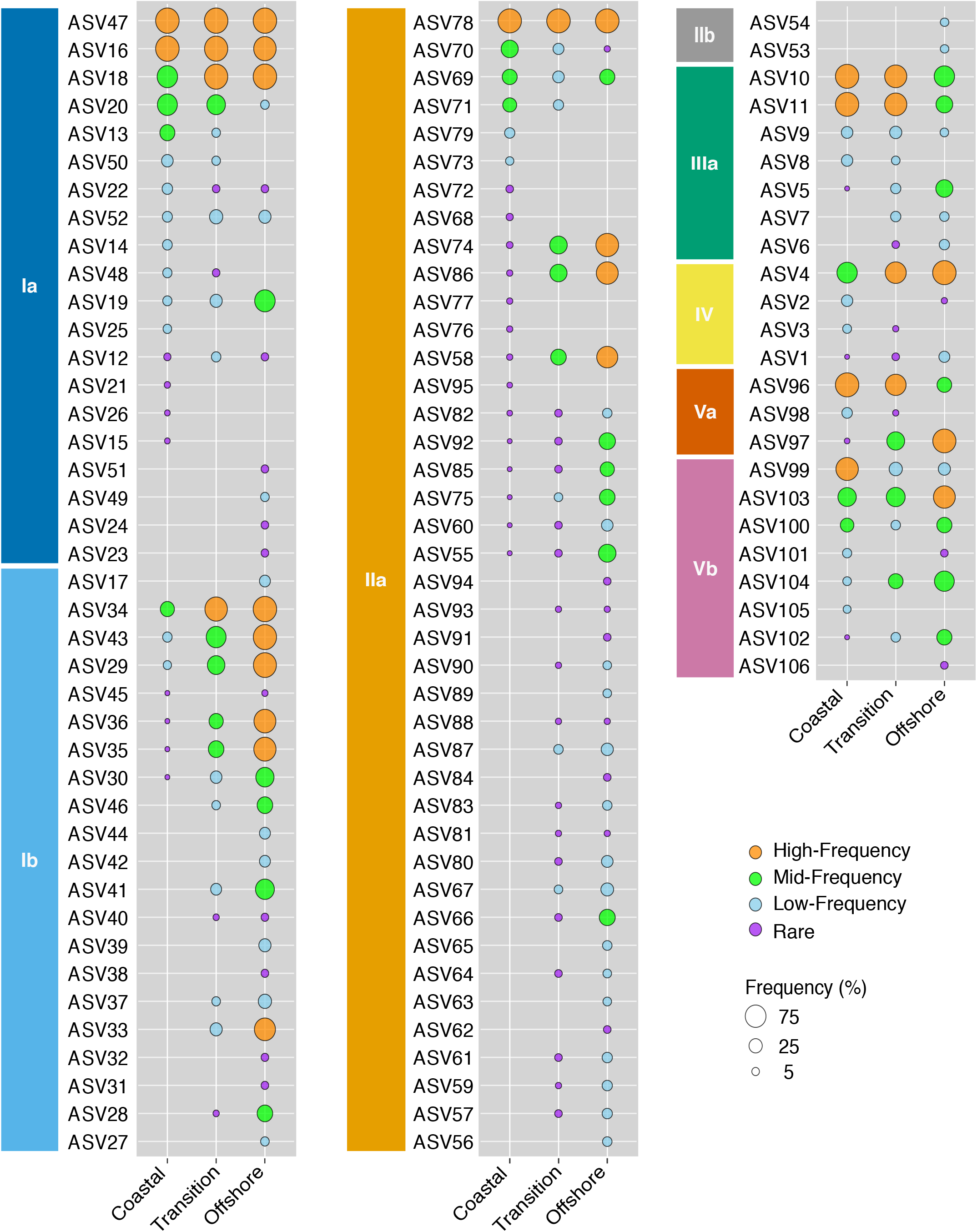
Frequency of detection of SAR11 ASVs in coastal (n=120), transition (n=40), and offshore (n=40) environments of KByT.

Rare ASVs were most commonly members of SAR11 subclade IIa (coastal n=14; transition n=16; offshore n=8). All subclades had an ASV that was high-frequency, with the exception of the exceedingly rare subclade IIb. Two high-frequency ASVs were ubiquitous across all 200 samples: ASV47 from subclade Ia and ASV78 from subclade IIa. Two high-frequency ASVs were ubiquitous across all coastal samples: ASV16 from subclade Ia and ASV96 from subclade Va. Three high-frequency ASVs were ubiquitous across all offshore samples: ASV18 from subclade Ia, ASV34 from Ib, and ASV4 from subclade IV.

In general, ubiquitous ASVs were also typically the most abundant. As a proportion of the total SAR11 fraction, ASV47 (28.5 ± 10.9%), ASV16 (24.2 ± 15.9%), ASV78 (8.5 ± 3.0%), ASV18 (6.9 ± 6.9%), ASV34 (3.9 ± 4.8%), and ASV96 (5.1 ± 4.2%) were the most abundant SAR11 ASVs across all samples (mean ± s.d.; n=200) (**Fig. 5)**. One exception was ASV4; at 0.8 ± 0.9% of the SAR11 fraction, it was only the 14th most abundant SAR11 ASV yet ubiquitous in the offshore. Within the coastal environment, the seven ubiquitous ASVs made up 83.0 ± 5.9% (mean ± s.d.; n=120) of the total SAR11 community, 77.9 ± 12.8% in the transition environment (n=40), and 62.8 ± 7.9% in the offshore (n=40) (**Fig. 5)**.

**Figure 5.**
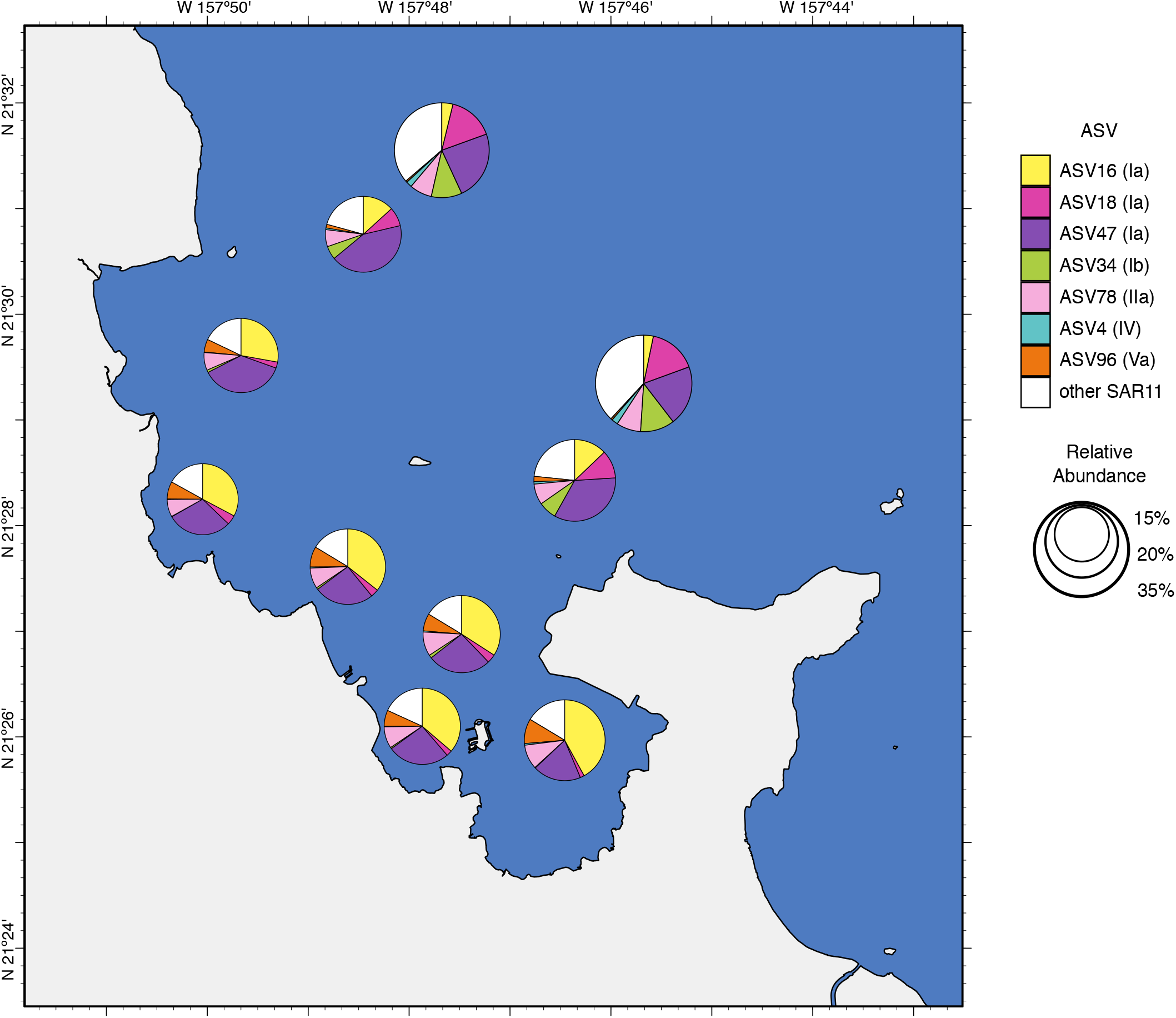
Average proportion of the total SAR11 relative abundance for the seven ASVs that are ubiquitous within coastal (ASV16, ASV96), offshore (ASV34, ASV18, ASV4), or both (ASV47, ASV78) environments. Circle size indicates the relative proportion of SAR11 in the total microbial community.

### Biogeography of SAR11 ASVs that match isolates

Of the 106 SAR11 ASVs detected in KByT, 11 were 100% identical to cultured strains of SAR11, including seven from subclade Ia (**Table 2**). This included the most abundant and ubiquitous ASV in this data set, ASV47 within subclade Ia, which was identical to the 16S rRNA gene of fifteen SAR11 strains including several isolated from Kāne‘ohe Bay (**Table 2; Table S4**). ASV16 and ASV18 from SAR11 subclade Ia each matched 100% to cultivated strains and possessed contrasting patterns of distribution, with ASV16 significantly more abundant in the coastal environment and ASV18 significantly more abundant in the transition and offshore environments (**Fig. 5; Table S3**). ASV78, the other ubiquitously-distributed ASV and the second-most abundant, was a 100% match to strain HIMB58 in subclade IIa. One subclade IIIa ASV (ASV11) with ubiquitous distribution in coastal samples was identical to strain HIMB114. Finally, ASV96 in subclade Va exactly matched strain HIMB59 and was ubiquitously distributed in the coastal environment.

**Table 2.** SAR11 ASVs recovered in KByT that match with 100% similarity to the 16S rRNA gene sequence of isolated strains.

| ASV | SAR11 subclade | Matching strains (Total no.) | Representative strain |
| --- | --- | --- | --- |
| ASV16<sup>a</sup> | Ia | 10 | HIMB140<sup>b</sup> |
| ASV47<sup>a</sup> | Ia | 15 | HIMB1412<sup>b</sup> |
| ASV18<sup>a</sup> | Ia | 10 | HIMB1437<sup>b</sup> |
| ASV22 | Ia | 1 | HIMB1463 |
| ASV20 | Ia | 1 | HIMB1427 |
| ASV12 | Ia | 1 | HIMB2304 |
| ASV50 | Ia | 1 | HIMB83 |
| ASV43<sup>a</sup> | Ib | 1 | RS40 |
| ASV78<sup>a</sup> | IIa | 1 | HIMB58 |
| ASV11<sup>a</sup> | IIIa | 1 | HIMB114 |
| ASV96<sup>a</sup> | Va | 1 | HIMB59 |

<sup>a</sup> Highly abundant within KByT.
<sup>b</sup> Full lists of strains appear in Table S4.

The majority (7 of 11) of ASVs with matches to previously isolated SAR11 strains occurred in high frequency and abundance. The remaining ASVs that matched isolated strains were all from subclade Ia (ASV50, ASV20, ASV22, ASV12), and were less frequent and abundant (**Table 2**).

## Discussion

The Hawaiian island landmasses provide a useful and convenient platform to investigate the ecotypic differentiation of planktonic marine bacteria over an abrupt environmental gradient. Across two years of monthly sampling, the coastal environment within Kāne‘ohe Bay, O‘ahu resolved itself as a marine *Synechococcus*-dominated system that contains elevated inorganic nutrients, chlorophyll *a*, and cellular abundances of planktonic marine bacteria. These features are consistent with an “Island Mass Effect”, where an increase in phytoplankton biomass proximate to near-island and atoll-reef ecosystems is caused by localized increases in nutrient delivery through physical oceanographic, biological, land-based, and anthropogenic processes [65]. A short, three nautical mile transit from the interior of the bay across a transition zone leads to a *Prochlorococcus*-dominated system characterized by depressed inorganic nutrients, low chlorophyll *a* concentrations, and lower abundances of planktonic marine bacteria-characteristics typical of offshore waters [66].

The persistent differences between coastal and offshore environments across the KByT transect also manifested in distinct distributions of SAR11 marine bacteria. While SAR11 accounted for roughly 20% of the microbial community within the coastal environment of Kāne‘ohe Bay, their relative abundance sharply increased to 30-35% in the offshore waters surrounding the bay. In fact, seven of eight SAR11 subclades detected throughout KByT displayed distinct patterns of distribution across coastal Kāne‘ohe Bay and the adjacent offshore, the most dramatic of which were an increase in abundance of subclades Ib and IIa in the offshore. Increases in these two subclades have been associated with oligotrophic ocean gyres [67] and decreasing salinity and nutrients [68]. Our results show that the environmental differences among coastal Kāne‘ohe Bay and the neighboring offshore is a strong determinant of SAR11 subclade distribution, providing further support that adaptations to environmental niches may shape the evolution of a majority of SAR11 marine bacteria [16].

In previous studies, abiotic parameters including depth, salinity, temperature, and latitude have been identified as primary drivers of the distribution of SAR11 subclades [8,11,16,23-26]. We observed a fundamental change in the picocyanobacteria that dominate coastal versus offshore environments across the KByT system, and hypothesize that this is a major driver determining the distinct distributions of SAR11 subclades and ASVs across KByT. Some evidence of how SAR11 populations may specialize in coastal versus oceanic environments comes from analysis of genomes sequenced from isolated SAR11 strains belonging to subclade Ia [69,70]. For example, genomic analyses of coastal, high-latitude subclade Ia strain HTCC1062 and ocean gyre, low-latitude subclade Ia strain HTCC7211 have shown that the coastal strain is capable of utilizing glucose for chemoheterotrophic growth, while the ocean gyre strain could not [69]. This appears to be a variable metabolic property within the SAR11 Ia subclade and is more commonly detected in productive environments that offer higher concentrations of labile carbon sources [69]. Recent genomic investigations and co-culture studies are expanding our understanding of the diverse phytoplankton byproducts SAR11 cells are able to metabolize, such as volatile organic compounds [71,72] and dimethyl arsenate [70], emphasizing the importance of co-evolutionary relationships between SAR11 and phytoplankton [73,74], and further suggesting that productivity plays a significant role in structuring SAR11 genetic and ecological diversity.

Across the three environments of KByT, the majority of SAR11 relative abundance was dominated by seven ASVs. This observation was similar to that by Ortmann & Santos [75], who found that the ten most abundant SAR11 OTUs represented > 80% of all of the SAR11 sequences collected from surface seawaters along a transect from coastal Mobile Bay, Alabama, to the offshore Gulf of Mexico. It is intriguing that the seven ASVs that dominated the SAR11 community spanned five of eight major subclades detected in this study. Given the evidence that some of these subclades appear to fulfill unique ecological niches [8,9,21,76], the reduction of interspecific competition and thus the capacity for coexistence among these dominant inter-subclade ASVs may be achieved through subtle resource-partitioning [77]. On the other hand, intense intraspecific competition may explain why most subclades are represented by a single highly-abundant ASV [78]. Subclade Ia had three ASVs (ASV16, 18, 47) within the seven most abundant, which is not surprising considering that it is the most abundant SAR11 subclade in surface oceans overall [21]. Our observation that ASV16 is more abundant nearshore and ASV18 is more abundant offshore provides evidence that their coexistence may in part derive from functional differences between closely related SAR11 ASVs that result in their differential distribution across end member environments of this study system.

In our study, spring and summer coincide with the dry season in Hawai‘i; a period of increased salinity, increased solar irradiance, and diminished dissolved organic nutrients in the coastal environment due in part to decreased nutrient delivery from freshwater streams [79,80]. At coastal KByT stations within Kāne‘ohe Bay, notable seasonal changes during spring including increased salinity, *Synechococcus* and heterotrophic bacteria cellular abundance, and an increase in the total relative abundance of SAR11 were observed. Based on our results and those of past studies [41,80], it appears that SAR11 and *Synechococcus* within Kāne‘ohe Bay tend to increase in abundance during dry periods and decrease seasonally and periodically (i.e. immediately following storm events) with increased rainfall and nutrient concentrations.

Seasonal changes in the diversity and abundance of individual SAR11 subclades have been observed in temperate [24,81], tropical [16], freshwater [82], and subtropical environments [83]. While significant seasonal changes in the alpha diversity of SAR11 subclades similar to the winter diversity minima reported by Salter and colleagues [24] were not observed in KByT, we did discern strong seasonal patterns in the relative abundance of SAR11 subclades and within individual ASVs. At the offshore stations of KByT, we found that SAR11 subclades Ia and Va increased during spring, IIIa peaked during fall, and IV peaked during winter in a similar fashion to observations in surface waters of the Sargasso Sea [16]. However, at the coastal stations of KByT, subclade Ia peaked in relative abundance during the winter while subclade IV peaked during spring. Several factors may contribute to differences in subclade seasonality between the coastal and offshore environments. First, the different ASVs that comprise these subclades in each environment may display heterogeneous responses to seasonal changes. Second, seasonal fluctuations in the structure and activity of the phytoplankton and cyanobacterial communities that make up the base of the food web that are specific to the coastal or offshore environment may elicit changes in the growth of SAR11 ASVs. In the coastal environment, the average relative abundances of SAR11 and *Synechococcus* reached their maximum during spring. Similarly, in the offshore environment SAR11 and *Prochlorococcus* both reached their maximum average relative abundances during the summer. It is plausible that the abundance of the major consumer of dissolved organic matter, SAR11, is linked to the abundance of dominant photoautotrophs *Synechococcus* and *Prochlorococcus* in the coastal and offshore environments, respectively [84].

Relatively little is known about the genetic structure of natural populations of marine microbes in coastal tropical environments, as most studies come from time-series in temperate systems [85,86] and oligotrophic ocean gyres [16,79,83,87]. Our time-series analyses from tropical coastal Kāne‘ohe Bay to the adjacent offshore system show distinct SAR11 communities across environments that are dominated by a few highly abundant ASVs. This study contributes to a growing knowledge of how coexisting, closely-related populations of marine bacteria are distributed across environmental gradients. Our observations of SAR11 marine bacteria at the scale of 16S rRNA gene ASVs, subclades, and total relative abundance provide a new understanding of how SAR11 genetic diversity partitions in distinct coastal and offshore systems.

## Acknowledgements

We thank Dr. Catherine M. Foley for her generous help with the creation of maps, Evelyn Hoffman, Helen Li, and Rachel Ouye for their laboratory and field assistance, Daniel Schar for assistance with fluorometric measurements, Jason Jones, Rebecca Weible, and Evan Barba for assistance with sample collection, Jed Fuhrman for providing mock microbial communities to use in the 16S rRNA gene analyses, and Brian Powell for his advice regarding the delineation of seasons. This research was supported by funding from the National Science Foundation (grant OCE-1538628 to MSR) and the National Science Foundation Graduate Research Fellowship Program to SJT. This is SOEST contribution xxx and HIMB contribution xxx.

## Supplementary Figure Legends

**Figure S1. A)** Harmonic mean function fit to sea surface water temperature from Kāne‘ohe Bay, Hawai‘i. **B)** Annual time-derivate for all 10 years of data.

**Figure S2**. Phylogenetic analysis of SAR11 16S rRNA gene ASVs recovered through KByT. Cultivated isolates are indicated by “str.” (strain). NCBI accession numbers of reference sequences are included in parentheses.

**Figure S3**. nMDS of a Bray-Curtis distance matrix calculated from temperature, salinity, pH, cellular abundances of heterotrophic bacteria, cyanobacteria, and eukaryotic picophytoplankton, and chlorophyll-a concentrations for the entire KByT dataset (n=200 samples). nMDS stress was 0.03.

**Figure S4**. Average relative abundance of the fourteen most abundant order-level 16S rRNA gene-based groups across coastal (n=120), transition (n=40), and offshore (n=40) environments sampled over two years of KByT. Taxonomy based on Silva v132.

**Figure S5**. Relative abundance of *Synehococcus* and *Prochlorococcus* ASVs across KByT. n=20 samples per station.

**Figure S6**. Average abundance and standard deviation of **A)** Chloroplast, **B)** *Synechococcus*, and **C)** *Prochlorococcus* sequences relative to the total community in each environment and season. Seasonal comparisons with significant seasonal differences as defined by DESeq2 normalization are noted: * if p<0.05, ** if p≤0.01, and *** if p≤ 0.001, while non-significant comparisons are not noted.

**Figure S7. A)** Average abundance and standard deviation **of** SAR11 relative to the total community in each environment. **B)** Average abundance and standard deviation **of** SAR11 relative to the total community in each season. Comparisons with significant seasonal or environmental differences as defined by DESeq2 normalization are noted: * if p<0.05, ** if p≤0.01, and *** if p≤ 0.001, while non-significant comparisons are not noted.

**Figure S8**. Average abundance and standard deviation **of** SAR11 subclades relative to total SAR11 abundance in each environment. Comparisons with significant environmental differences as defined by DESeq2 normalization are noted: * if p<0.05, ** if p≤0.01, and *** if p≤ 0.001, while non-significant comparisons are not noted.

**Figure S9**. Average abundance and standard deviation of SAR11 subclades relative to total SAR11 abundance in each environment (coastal, transition, offshore) per season. Seasonal comparisons with significant seasonal differences as defined by DESeq2 normalization are noted: * if p<0.05, ** if p≤0.01, and *** if p≤ 0.001, while non-significant comparisons are not noted.

**Figure S10**. Heatmap indicating the relative abundance of SAR11 ASVs (rows) per sample (columns). ASVs are ordered vertically by subclade, and samples are ordered horizontally by environment, season, date sampled, and site. “E” next to ASVs denotes a DESeq2 significant difference between environments, while “S” denotes DESeq2 significant differences between seasons.

## Supplementary Tables Legends

**Table S1**. Environmental parameters and cellular abundances from KByT. Samples are averaged (mean ± s.d.) over environmental category and season.

**Table S2**. Comparisons of water column parameters and cellular counts over time and space using one-way ANOVA. Significance (Holm corrected p-values) are shown.

**Table S3:** Summary of SAR11 ASVs recovered from KByT. ASVs with significant DESeq2 normalized differences across environments and seasons within each environment (C=coastal, T=transition, and O=offshore) are noted.

**Table S4**. Summary of KByT ASVs that match isolated SAR11 strains.

