## Supplemental Figures and Table S4 for "Spatial and temporal dynamics of SAR11 marine bacteria sampled across a nearshore to offshore transect in the tropical Pacific Ocean"

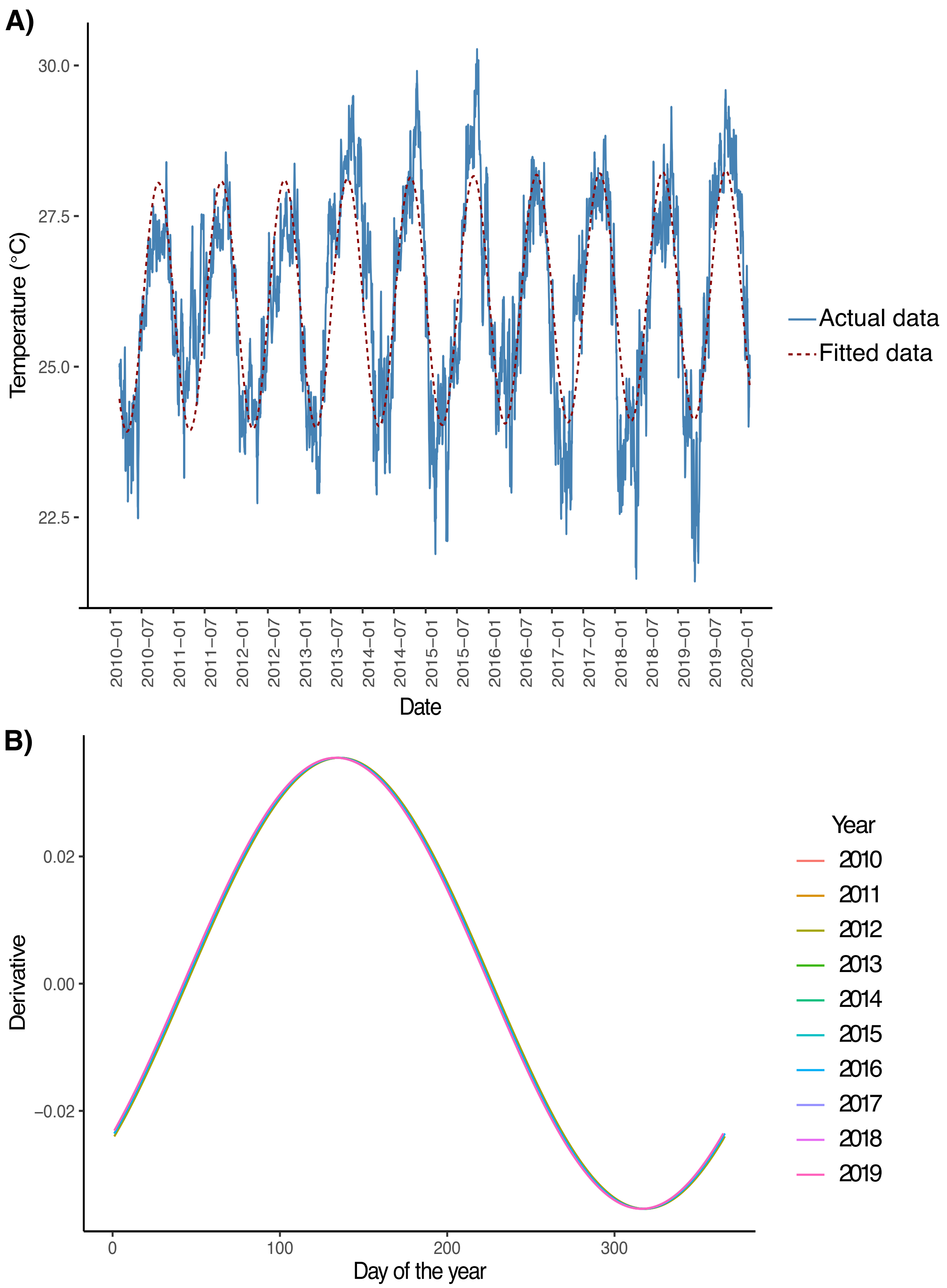
A)
30.0
27.5
25.0
22.5
Temperature (°C)
2010−01
2010−07
2011−01
2011−07
2012−01
2012−07
2013−01
2013−07
2014−01
2014−07
2015−01
2015−07
2016−01
2016−07
2017−01
2017−07
2018−01
2018−07
2019−01
2019−07
2020−01
Date
Actual data
Fitted data
B)
0.02
0.00
−0.02
Derivative
0
100
200
300
Day of the year
Year
2010
2011
2012
2013
2014
2015
2016
2017
2018
2019

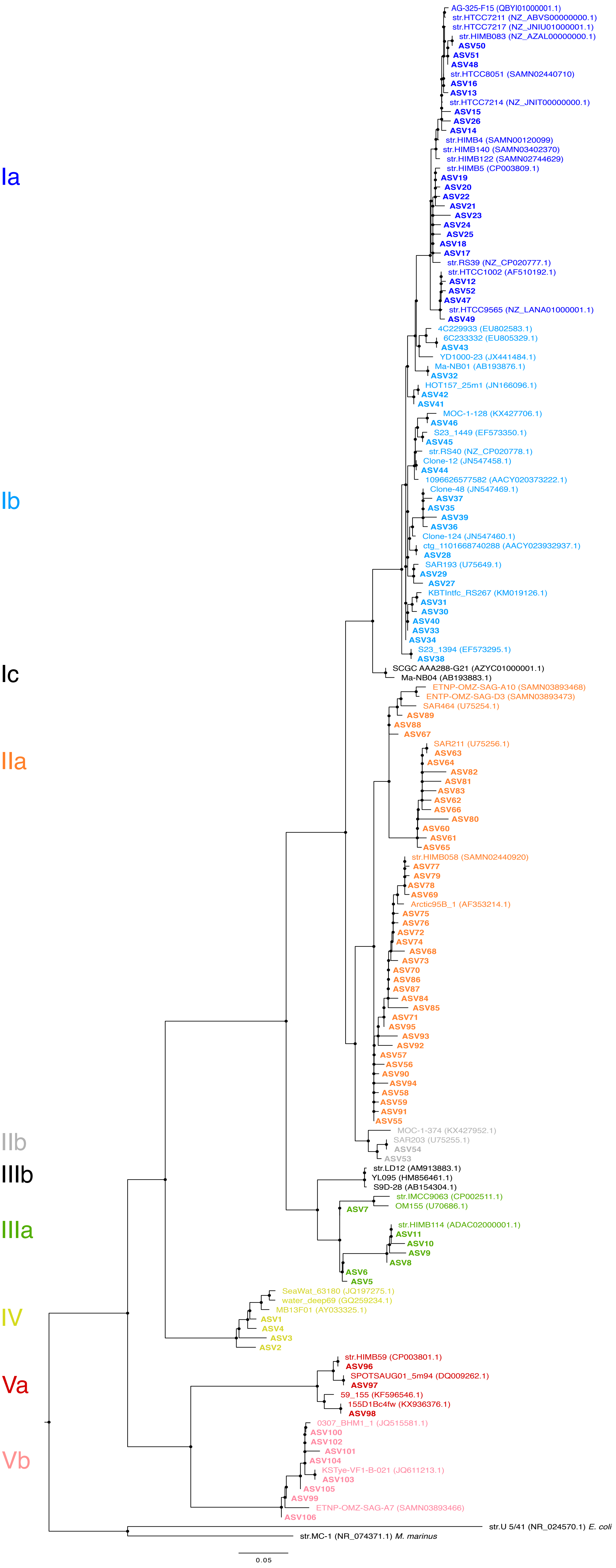

Ia
Ib
Ic
IIa
IIb
IIIb
IIIa
IV
Va
Vb
AG-325-F15 (QBYI01000001.1)
str.HTCC7211 (NZ_ABVS00000000.1)
str.HTCC7217 (NZ_JNIU01000001.1)
str.HIMB083 (NZ_AZAL00000000.1)
ASV50
ASV51
ASV48
str.HTCC8051 (SAMN02440710)
ASV16
ASV13
str.HTCC7214 (NZ_JNIT00000000.1)
ASV15
ASV26
ASV14
str.HIMB4 (SAMN00120099)
str.HIMB140 (SAMN03402370)
str.HIMB122 (SAMN02744629)
str.HIMB5 (CP003809.1)
ASV19
ASV20
ASV22
ASV21
ASV23
ASV24
ASV25
ASV18
ASV17
str.RS39 (NZ_CP020777.1)
str.HTCC1002 (AF510192.1)
ASV12
ASV52
ASV47
str.HTCC9565 (NZ_LANA01000001.1)
ASV49
4C229933 (EU802583.1)
6C233332 (EU805329.1)
ASV43
YD1000-23 (JX441484.1)
Ma-NB01 (AB193876.1)
ASV32
HOT157_25m1 (JN166096.1)
ASV42
ASV41
MOC-1-128 (KX427706.1)
ASV46
S23_1449 (EF573350.1)
ASV45
str.RS40 (NZ_CP020778.1)
Clone-12 (JN547458.1)
ASV44
1096626577582 (AACY020373222.1)
Clone-48 (JN547469.1)
ASV37
ASV35
ASV39
ASV36
Clone-124 (JN547460.1)
ctg_1101668740288 (AACY023932937.1)
ASV28
SAR193 (U75649.1)
ASV29
ASV27
KBTIntfc_RS267 (KM019126.1)
ASV31
ASV30
ASV40
ASV33
ASV34
S23_1394 (EF573295.1)
ASV38
SCGC AAA288-G21 (AZYC01000001.1)
Ma-NB04 (AB193883.1)
ETNP-OMZ-SAG-A10 (SAMN03893468)
ENTP-OMZ-SAG-D3 (SAMN03893473)
SAR464 (U75254.1)
ASV89
ASV88
ASV67
SAR211 (U75256.1)
ASV63
ASV64
ASV82
ASV81
ASV83
ASV62
ASV66
ASV80
ASV60
ASV61
ASV65
str.HIMB058 (SAMN02440920)
ASV77
ASV79
ASV78
ASV69
Arctic95B_1 (AF353214.1)
ASV75
ASV76
ASV72
ASV74
ASV68
ASV73
ASV70
ASV86
ASV87
ASV84
ASV85
ASV71
ASV95
ASV93
ASV92
ASV57
ASV56
ASV90
ASV94
ASV58
ASV59
ASV91
ASV55
MOC-1-374 (KX427952.1)
SAR203 (U75255.1)
ASV54
ASV53
str.LD12 (AM913883.1)
YL095 (HM856461.1)
S9D-28 (AB154304.1)
str.IMCC9063 (CP002511.1)
OM155 (U70686.1)
ASV7
str.HIMB114 (ADAC02000001.1)
ASV11
ASV10
ASV9
ASV8
ASV6
ASV5
SeaWat_63180 (JQ197275.1)
water_deep69 (GQ259234.1)
MB13F01 (AY033325.1)
ASV1
ASV4
ASV3
ASV2
str.HIMB59 (CP003801.1)
ASV96
SPOTSAUG01_5m94 (DQ009262.1)
ASV97
59_155 (KF596546.1)
155D1Bc4fw (KX936376.1)
ASV98
0307_BHM1_1 (JQ515581.1)
ASV100
ASV102
ASV101
ASV104
KSTye-VF1-B-021 (JQ611213.1)
ASV103
ASV105
ASV99
ETNP-OMZ-SAG-A7 (SAMN03893466)
ASV106
str.U 5/41 (NR_024570.1) E. coli
str.MC-1 (NR_074371.1) M. marinus
0.05

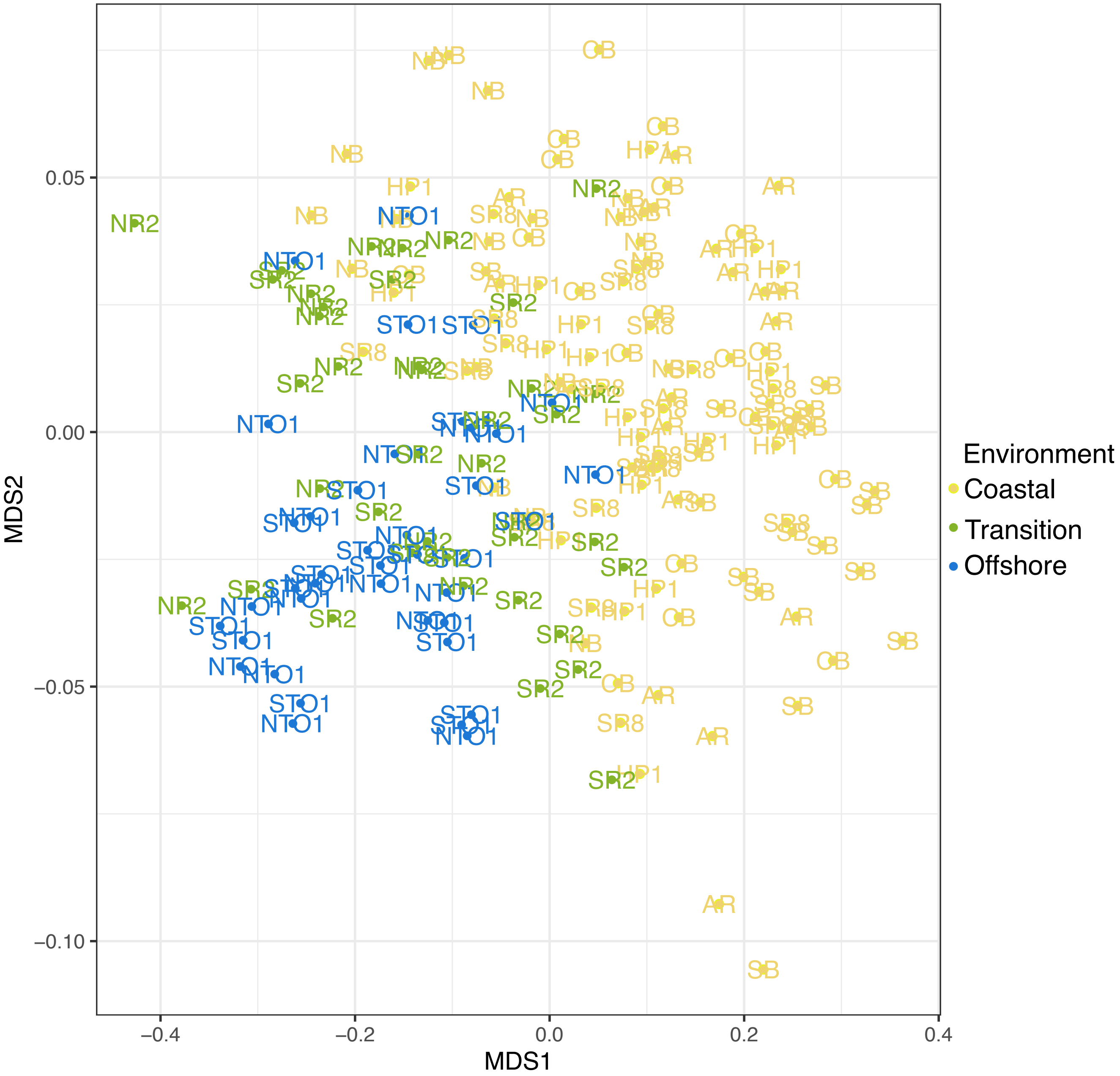

MDS2
0.05
0.00
−0.05
−0.10
MDS1
−0.4
−0.2
0.0
0.2
0.4
Environment
Coastal
Transition
Offshore

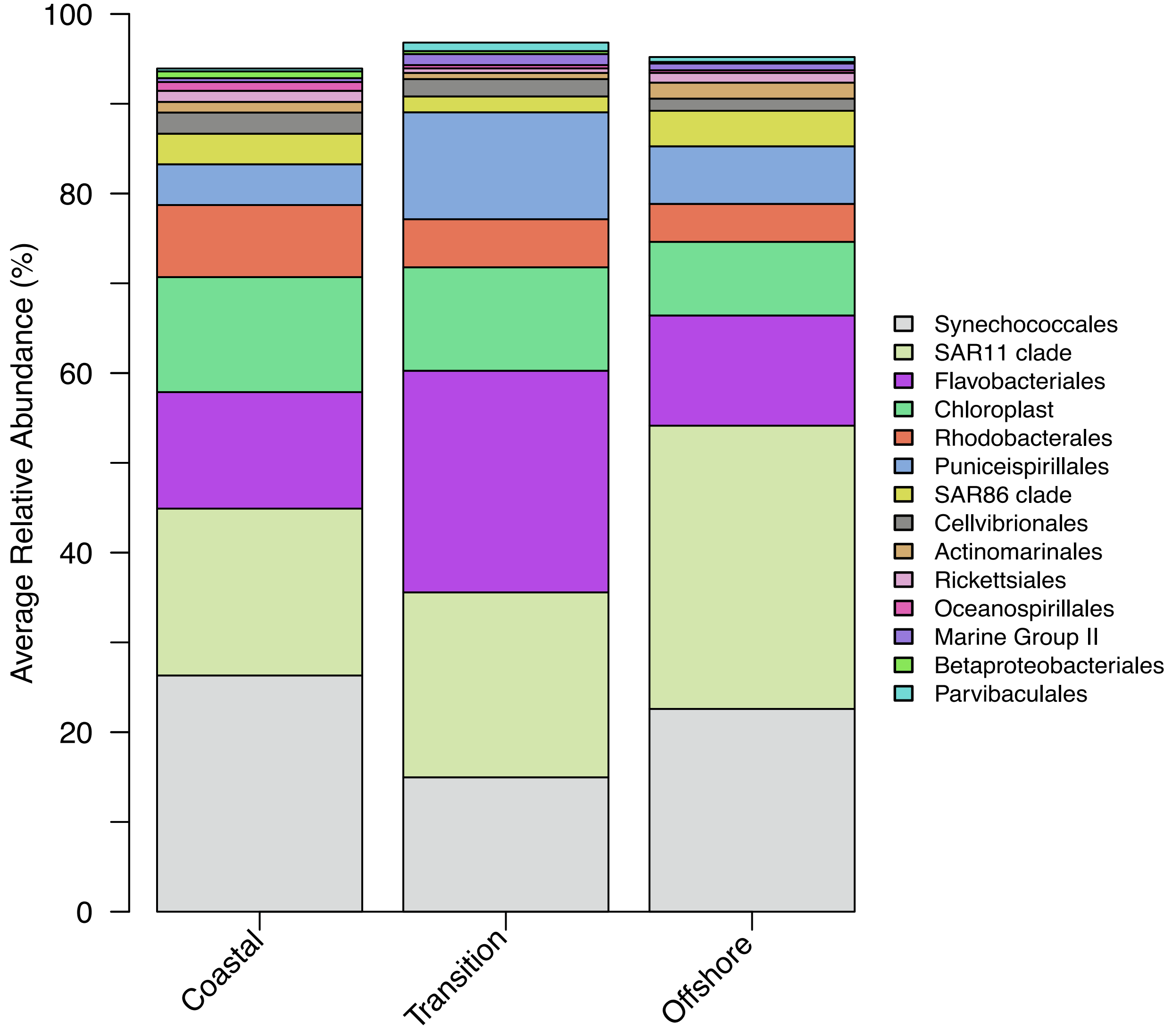

Average Relative Abundance (%)
100
80
60
40
20
0
Coastal
Transition
Offshore
Synechococcales
SAR11 clade
Flavobacteriales
Chloroplast
Rhodobacterales
Puniceispirillales
SAR86 clade
Cellvibrionales
Actinomarinales
Rickettsiales
Oceanospirillales
Marine Group II
Betaproteobacteriales
Parvibaculales

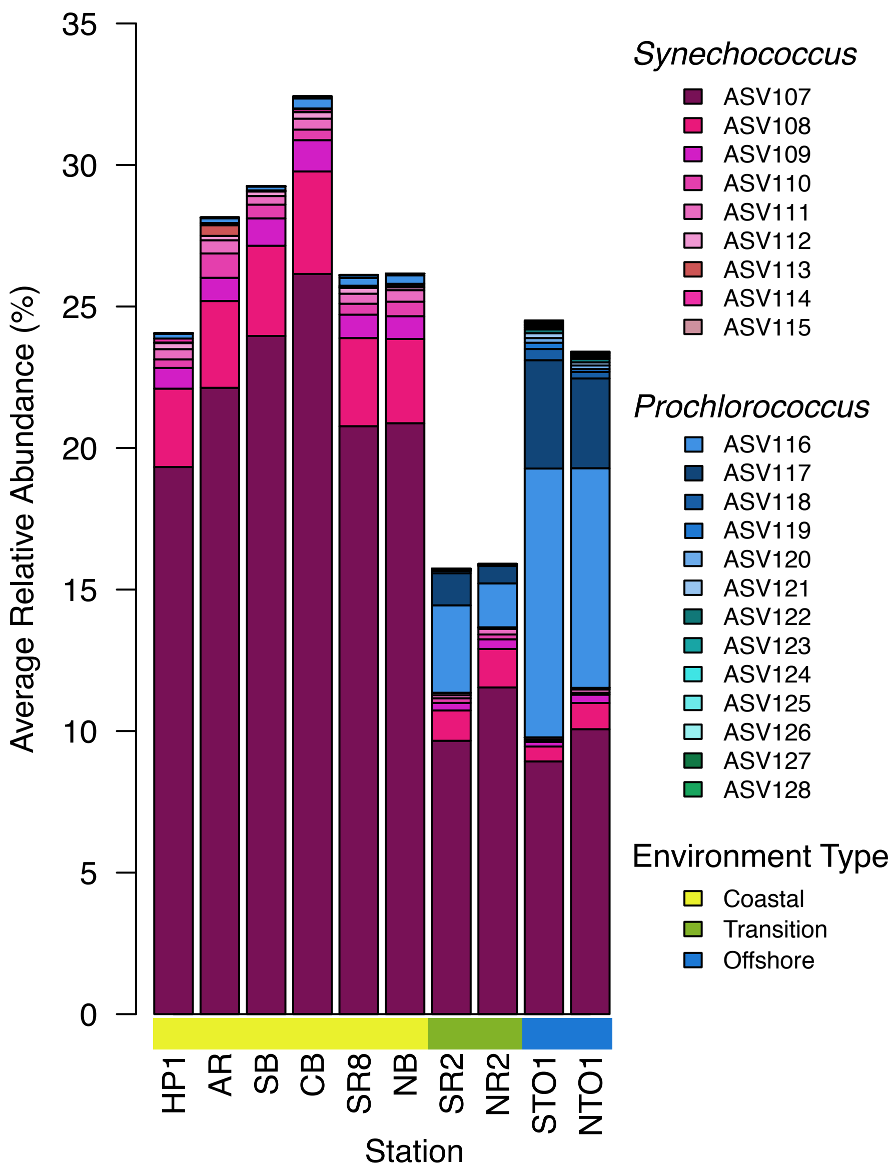

35
30
25
20
15
10
5
0
Average Relative Abundance (%)
HP1
AR
SB
CB
SR8
NB
SR2
NR2
STO1
NTO1
Station
Synechococcus
ASV107
ASV108
ASV109
ASV110
ASV111
ASV112
ASV113
ASV114
ASV115
Prochlorococcus
ASV116
ASV117
ASV118
ASV119
ASV120
ASV121
ASV122
ASV123
ASV124
ASV125
ASV126
ASV127
ASV128
Environment Type
Coastal
Transition
Offshore

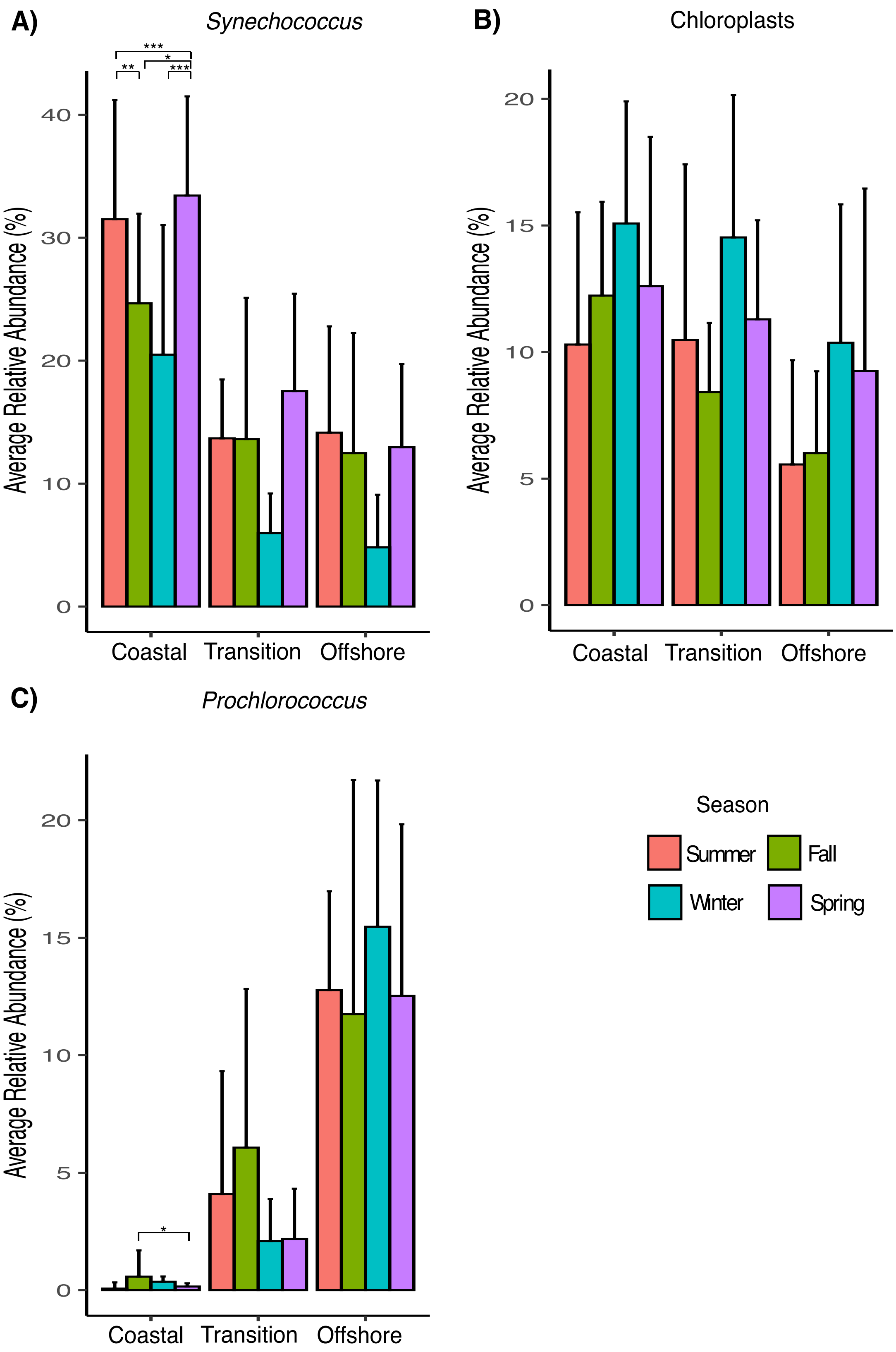

A)
Synechococcus
B)
Chloroplasts
C)
Prochlorococcus
Average Relative Abundance (%)
Coastal
Transition
Offshore
Season
Summer
Fall
Winter
Spring

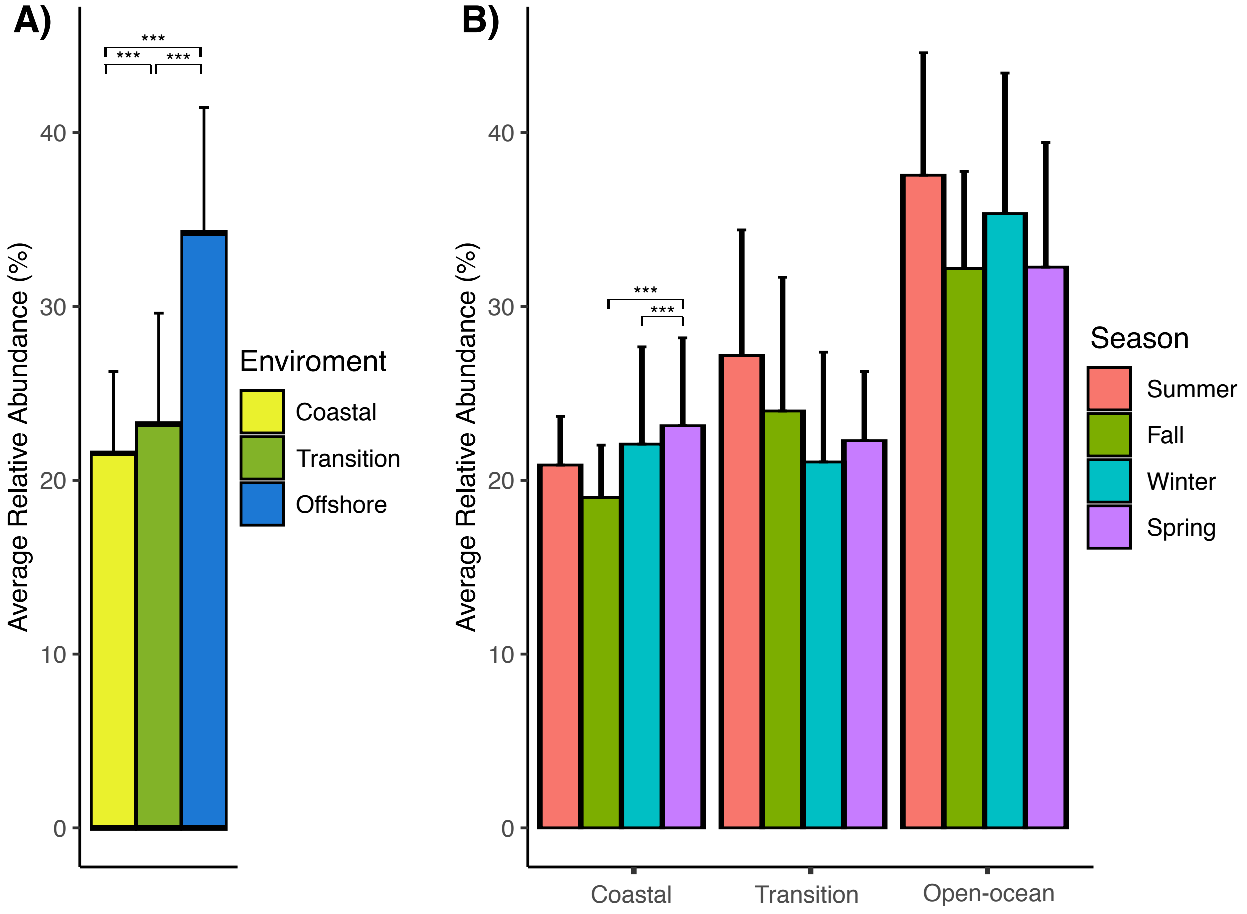
A)
B)
Average Relative Abundance (%)
0
10
20
30
40
***
***
***
Enviroment
Coastal
Transition
Offshore
Season
Summer
Fall
Winter
Spring
Coastal
Transition
Open-ocean

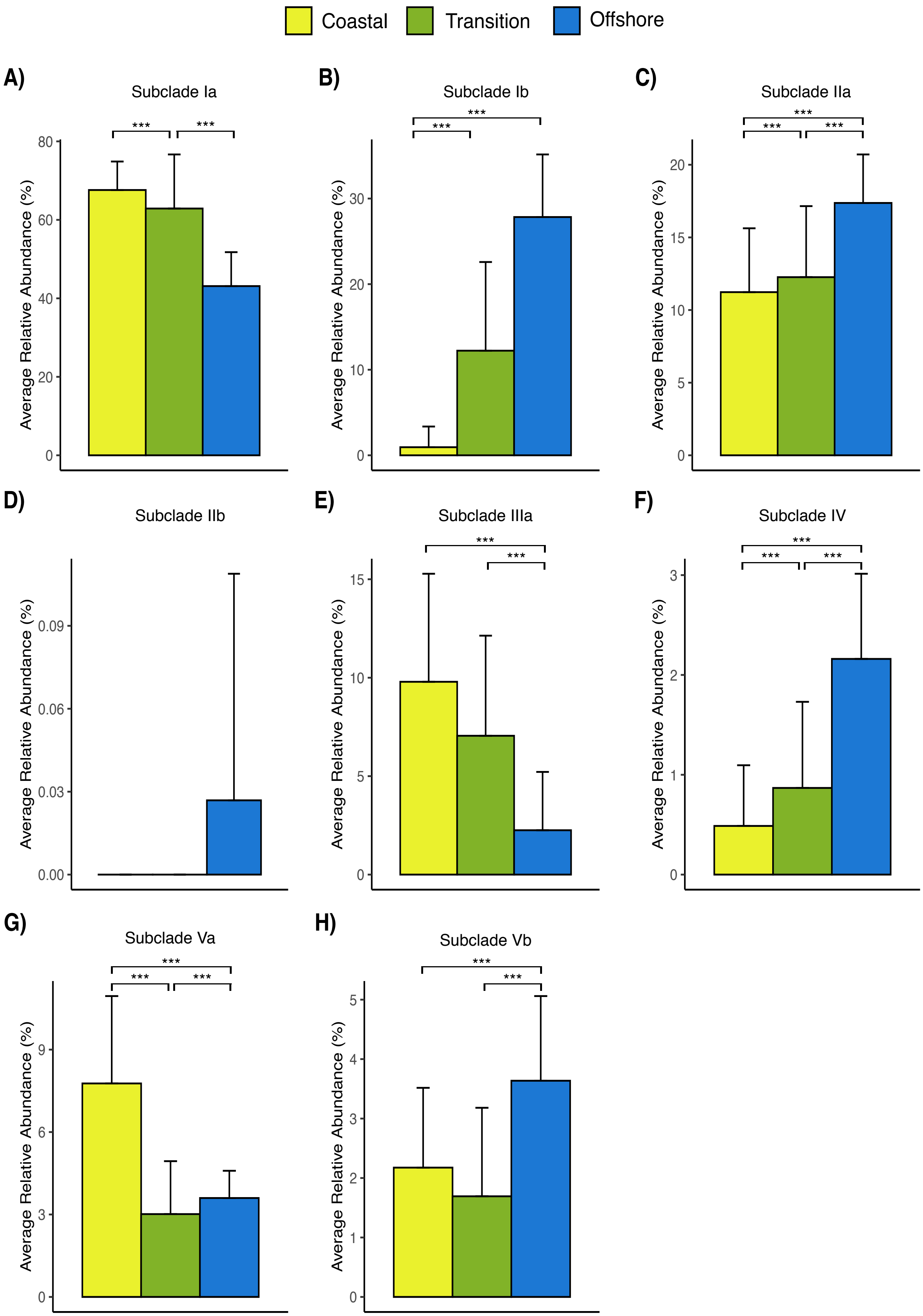
Coastal
Transition
Offshore
A)
Subclade Ia
B)
Subclade Ib
C)
Subclade IIa
D)
Subclade IIb
E)
Subclade IIIa
F)
Subclade IV
G)
Subclade Va
H)
Subclade Vb
Average Relative Abundance (%)
***

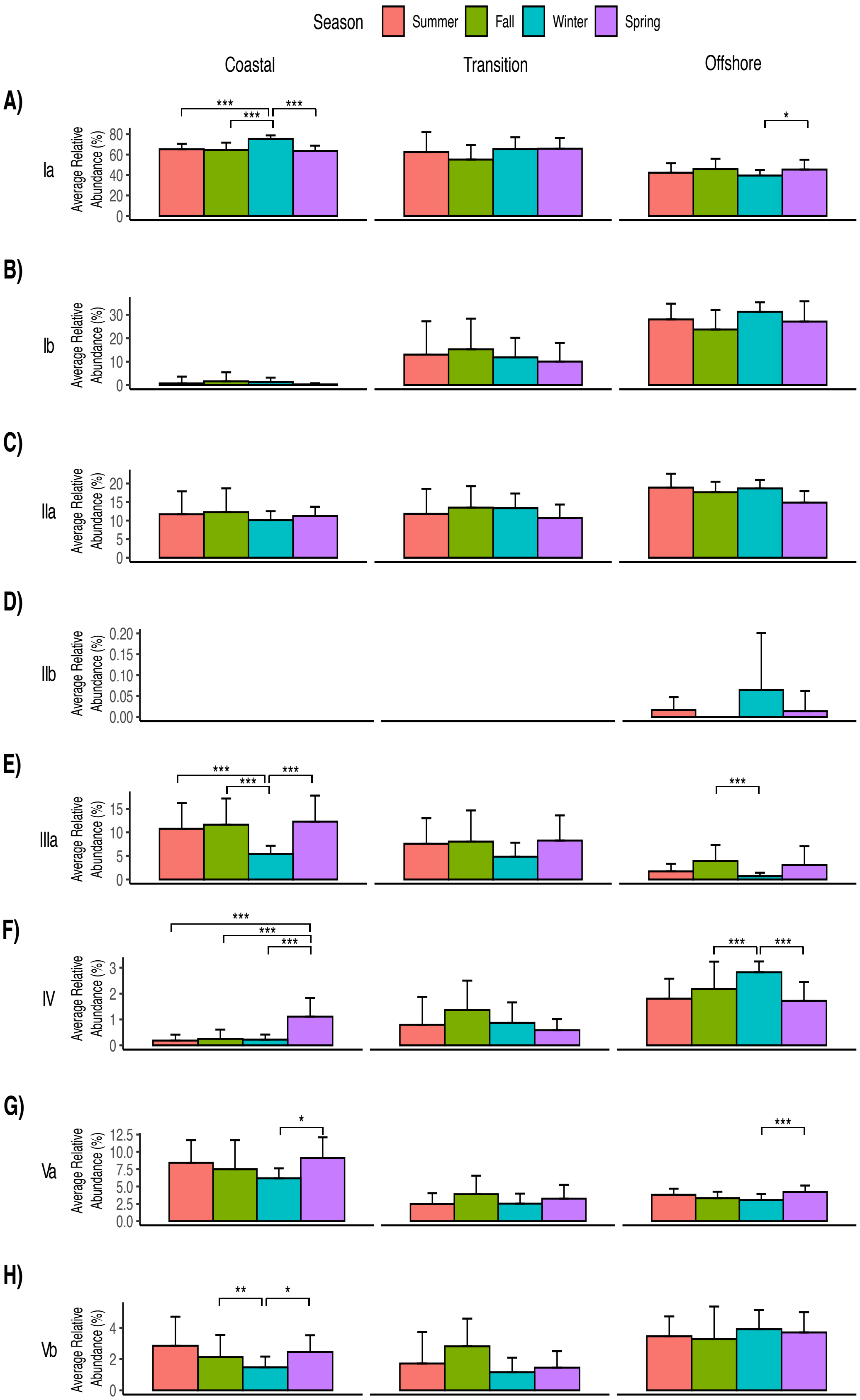

Season
Summer
Fall
Winter
Spring
Coastal
Transition
Offshore
A)
Ia
B)
Ib
C)
IIa
D)
IIb
E)
IIIa
F)
IV
G)
Va
H)
Vb
Average Relative Abundance (%)

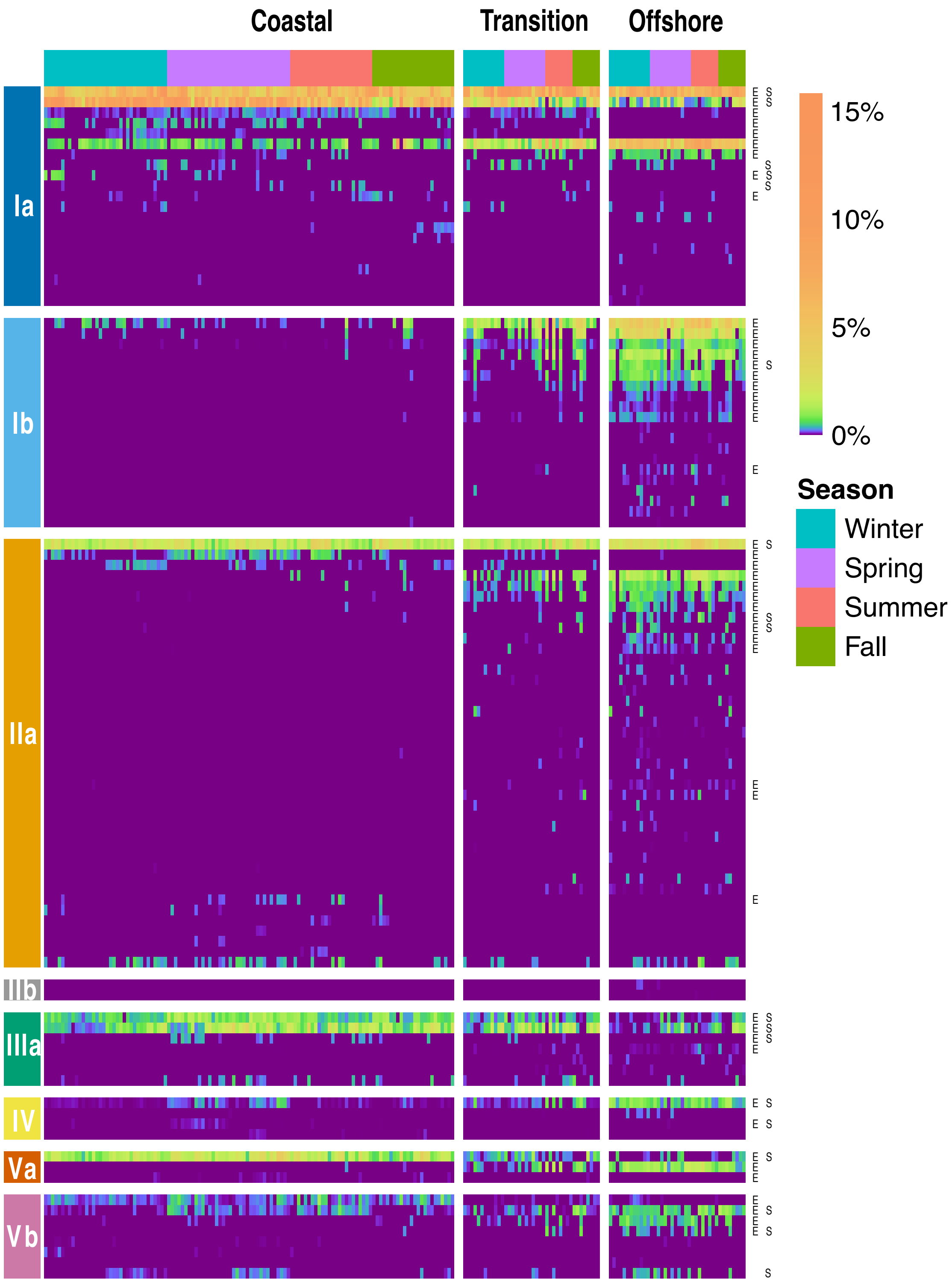
Coastal
Transition
Offshore
Ia
Ib
IIa
IIb
IIIa
IV
Va
Vb
15%
10%
5%
0%
Season
Winter
Spring
Summer
Fall

**Table S4.** Summary of KByT ASVs that match isolated SAR11 strains.

| References | ASV | Subclade |
|---|---|---|
| HTCC1062, HTCC1002, HTCC9565, RS39, HTCC1016, HTCC1013, HTCC1040, HIMB1412, HIMB1420, HIMB1436, HIMB1444, HIMB1456, HIMB2201, HIMB2247, HIMB2250 | ASV47* | Ia |
| HIMB140, HTCC7211, HTCC7214, HTCC8051, HIMB4, HTCC7217, HIMB1321, HTCC9022, HIMB1430c, HIMB2211 | ASV16* | Ia |
| HIMB1402, HIMB1437, HIMB1835, HIMB1838, HIMB1863, HIMB2187, HIMB2200, HIMB2204, HIMB2215, HIMB2226 | ASV18* | Ia |
| HIMB1427 | ASV20 | Ia |
| HIMB83 | ASV50 | Ia |
| HIMB1463 | ASV22 | Ia |
| HIMB2304 | ASV12 | Ia |
| RS40 | ASV43* | Ib |
| HIMB58 | ASV78* | IIa |
| HIMB114 | ASV11* | IIIa |
| HIMB59 | ASV96* | Va |

[a]ASVs with asterisks (*) represent ASVs that are highly abundant within our dataset.
